# Ultrastructural dynamics of basal bodies during microgamete formation and fertilisation in *Plasmodium*

**DOI:** 10.64898/2026.09.06.749571

**Authors:** Molly Hair, Ryuji Yanase, David J P Ferguson, Declan Brady, Richard Wheeler, Rita Tewari, Sue Vaughan

## Abstract

Cilia and flagella are microtubule-based organelles found in a wide range of eukaryotic organisms that form cilia and flagella are assembled from basal bodies. In the malaria parasite, eight flagellated microgametes are assembled from 8 basal bodies that form *de novo* as a single group (not next to a parent basal body) in the cytoplasm of microgametocytes extremely rapidly, in as little 8 minutes but is not synchronised. The flagellated microgametes exit from a microgametocyte, each consisting of an axoneme and a haploid nucleus. Fertilisation occurs via a *HAP2*-mediated fusion with the macrogamete. Despite the essential role of microgametes in malaria transmission and being the only flagellated stage, little is known about the process of basal body formation, biogenesis at the ultrastructure level and their role in fertilisation. We used dual axis serial section electron tomography (ssET) to reveal the unusual single microtubule structure of the basal body and discovered an associated electron dense basal body granule. Using whole cell reconstructions of microgametocytes, free microgametes and macrogametes by serial block face scanning electron microscopy (SBF-SEM), we discovered a deuterosome-like structure only present during the initial formation of 8 basal bodies that could be a nucleating platform for *de novo* basal body formation. Finally, we reveal that entry of the microgamete into the macrogamete occurs via directed event at a single point of entry with the basal body and granule leading entry. These discoveries highlight the essential functions of basal bodies from initial microgamete assembly to male-female gamete fertilisation.

## Introduction

Basal bodies are specialised centrioles and are microtubule-based barrel-shaped organelles usually composed of a 9 + 0 triplet arrangement of microtubules (A-B-C tubules) that nucleate the formation of flagella and cilia in eukaryotic organisms. The axoneme of flagella is composed on a 9 + 2 microtubule arrangement, with 9 outer double microtubules (A-B tubules) and 2 central pair microtubules. Cells often maintain a constant number of basal bodies by templated duplication from an existing basal body prior to cell division. Some cells can quickly produce many basal bodies *de novo,* such as during ciliogenesis to produce thousands of cilia such as in terminally differentiated mammalian epithelial cells which nucleate basal bodies at a deuterosome to produce hundreds of cilia (Briggs *et al*., 2004; Dawe, Farr and Gull, 2007; Hodges *et al*., 2010; Carvalho-Santos *et al*., 2011). This was first discovered as an electron dense structure by thin section transmission electron microscopy by Steinman (Steinman, 1968) and is only present in the early stages of basal body assembly and disappears following basal body assembly (Zhao *et al*., 2013).

Most *Plasmodium* life cycle stages lack centrioles or basal bodies, and basal bodies are only present during the sexual stages of the life cycle in *Plasmodium species* in the mosquito midgut (Sinden, 1991; Francia, Dubremetz and Morrissette, 2016). Environmental cues in the mosquito midgut trigger activation of the male gametocyte (microgametocyte) to undergo three rounds of intranuclear mitosis and assemble 8 basal bodies and axonemes in the cytoplasm without the need for intraflagellar transport (IFT) (Marques *et al*., 2015; Francia, Dubremetz and Morrissette, 2016; Zeeshan *et al*., 2022). This process is preparation for production of eight haploid male gametes (microgametes) that exflagellate from the microgametocyte and fertilise the female gamete (macrogamete) (Sinden, 1983; Guttery, Holder and Tewari, 2012; Dash *et al*., 2022).

There is a core set of highly conserved proteins essential for basal body assembly in most eukaryotic organisms and *Plasmodium* has homologs of at least three of these proteins (SAS6, SAS4 and BLD10/CEP135) (Hodges *et al*., 2010; Carvalho-Santos *et al*., 2011; Talman *et al*., 2014; Marques *et al*., 2015; Zeeshan *et al*., 2022). This suggests that much of the molecular machinery is conserved, but detailed studies of both the structural architecture and function of basal bodies in *Plasmodium species* are not well understood. For example, there are only rare thin section electron microscopy images showing basal body substructure with singlet microtubules (Sinden, Canning and Spain, 1976; Francia, Dubremetz and Morrissette, 2016; Zeeshan *et al*., 2022; Yang *et al*., 2025).

During exflagellation, each microgamete exits the microgametocyte with the basal body leading the exit, suggesting that basal bodies could also play a role in exflagellation, but mechanisms involved in exflagellation are not currently understood. Following exflagellation, microgametes and macrogametes undergo gamete-to-gamete fusion via a *HAP2* fusogen, but the precise mechanism of gamete fusion or how the haploid genome is delivered into the macrogamete is also not well understood (Liu *et al*., 2008; Feng *et al*., 2021; Kumar *et al*., 2022; Russell *et al*., 2023).

Gametogenesis is a vital step in the *Plasmodium* life cycle and there is research on this stage for the development of potential transmission blocking vaccines (Williamson *et al*., 1996; Patel and Tolia, 2021; Shasha *et al*., 2022). Little is known about the detailed structural architecture and biogenesis of the basal bodies within the microgametocyte and what role, if any, they have in microgamete function during fertilisation. Here, we have used dual axis serial section electron tomography (ssET) to investigate the detail structural architecture of basal bodies. We reveal that the *P. berghei* basal body is composed of an unusually short ∼50 nm nine singlet microtubule barrel instead of the usual triplet arrangement and has an electron dense granule associated with the proximal end of the basal body. To investigate basal body biogenesis and fertilisation we used serial block face scanning electron microscopy (SBF-SEM) to reconstruct whole individual microgametocytes, free microgametes and macrogametes. We have discovered a deuterosome-like structure that may act as a nucleating platform during the initial formation of 8 basal bodies. Finally, we discovered that entry of the microgamete occurs via directed event at a single point of entry with the basal body and granule leading the entry and fusion events occurring from the proximal end of the microgamete towards the distal end during fertilisation. This work provides novel findings in basal body ultrastructure during rapid gamete formation as well as its association in fertilisations events in *Plasmodium berghei*.

## Results

### *Plasmodium berghei* basal bodies are composed of a short single microtubule basal body and an associated basal body granule

Preceding stages of the life cycle (asexual proliferation) lack a centriole/basal body, therefore basal bodies must be formed *de novo* and rapidly during microgamete formation. We investigated the 3D ultrastructure of basal bodies to gain a clearer understanding of similarities and differences to other eukaryotic organisms. Serial dual axis serial tomograph (ssET) was used to examine complete microgametocyte basal bodies and the proximal part of the 9 + 2 microtubule axoneme of the flagellum (n=40 serial dual axis tomograms). These revealed an unusual basal body architecture when compared to other eukaryotic organisms. There was a very short basal body of 50 nm in length consisting of a 9+0 singlet microtubule arrangement embedded in electron dense material (Figure 1A, Movie 1). There was a typical 9 + 2 doublet microtubule axoneme like other eukaryotes, but one of the 2 central pair microtubules assembled more proximal than the other, as shown in *P. yoelii* (Yang *et al*., 2025) (Figure 1A - central pair microtubules, Movie 1).

**Figure 1.**
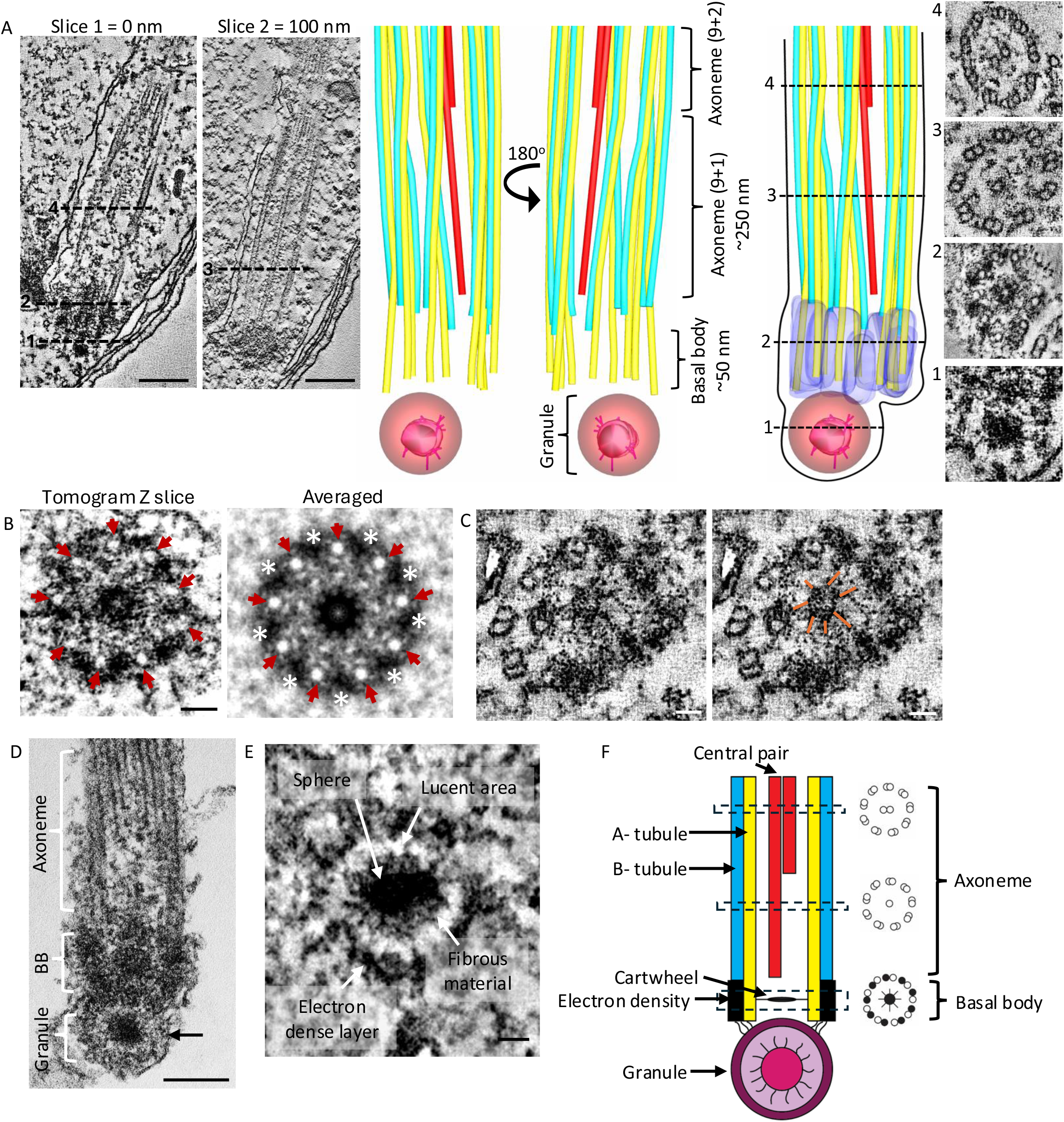
*Plasmodium berghei* basal body is composed of a single microtubule basal body and associated granule. **A:** Tomogram slices and segmentation showing the ultrastructure of a basal body and axoneme within a microgametocyte. Segmentation shows the granule (composed of a central electron density – pink, filamentous material – dark pink, and an electron dense ring – red, see below), the basal body is formed of 9 single microtubules (yellow) which are each associated with an electron density (purple), a second outer doublet microtubule (B-tubule blue) forms on the side of the single A-tubule distal to the basal body to form the axoneme. One central pair microtubule (red) of the axoneme begins assembly ∼250nm before the second central pair microtubule to form the 9 + 2 axoneme of the flagellum. Tomogram slices through the (1) granule (2) basal body, (3) axoneme with one central pair present and (4) a both central pairs present. 1-3 correspond to the tomogram slices through the basal body and axoneme. Scale bar = 200 nm. **B:** Tomogram slice of a basal body showing nine singlet microtubules (red arrows). Scale bar = 50 nm. Ninefold rotational averaging of the basal body structure of tomogram slice highlighting nine singlet microtubules (red arrows) and nine associated electron densities (white asterisks – shown in A in purple). **C:** Tomogram slice of basal body cartwheel (orange lines) within the basal body. Scale bar = 50 nm. **D:** Transmission electron micrograph (TEM) of the basal body associated granule (arrow) proximal to the basal body and axoneme. Scale bar = 100 nm. **E:** Tomogram slice of the basal body associated granule, highlighting the structural elements of the granule. Scale bar = 50 nm. **F:** Cartoon summary showing the basal body associated granule proximal to the basal body which is composed of nine singlet alpha microtubules and nine electron densities. Distal to the basal body, the axoneme displays a 9+1 configuration, with one central pair microtubule positioned more proximal than the other, transitioning into a classical 9+2 axoneme.

Electron dense material (densities) surrounded each of the nine single microtubules of the basal body, so ninefold rotational averaging was used to investigate the precise architecture of the A-tubule and electron densities. This method revealed nine singlet microtubules with nine electron dense masses positioned between each singlet microtubule (Figure 1B; singlet microtubules – arrows, electron dense masses - stars). These electron densities between each of the single microtubules extended almost the length of the 50nm basal body (Figure 1A – red arrow and the dark blue structure shown in the 3D reconstruction in Figure 1A). Eukaryotic basal bodies contain a cartwheel scaffolding structure at the proximal end projecting outward towards the nine triplet microtubules (Nakazawa *et al*., 2007; Meehl *et al*., 2016; Kantsadi *et al*., 2022) and we were able to resolve a cartwheel structure at the proximal end of basal bodies. (Figure 1C - orange lines).

There was a distinct electron dense, approximately spherical structure, associated with the proximal end of all basal bodies (Figure 1A – 1 - granule, 1D – arrow; n-40 tomograms). This structure was previously observed in thin section transmission electron microscopy (TEM) and was called a juxta-kinetosomal granule in *P. yoelii nigeriensis* (Sinden, Canning and Spain, 1976) but no more recent studies have been published. The kinetosome is a former term for a basal body, so we have re-named the structure a basal body-associated granule. Our datasets revealed the basal body-associated granule was composed of three distinct layers; a central electron-dense sphere, electron-lucent area surrounding the sphere and a further electron dense outer layer (Figure 1A – 1, 1E). The diameter of the granule was ∼150 nm, with the central sphere measuring ∼50 nm in diameter (n=40). Within the electron-lucent layer there was fibrous material that likely connected the central sphere to the outer electron dense layer (Figure 1E – fibrous material). Furthermore, the granule appeared to connect directly to the proximal end of the basal body (Figure 1D). This architecture is summarised in Figure 1F.

### Basal body biogenesis initiates with *de novo* formation of 8 basal bodies surrounding a deuterosome-like platform

The discovery of this unusual basal body ultrastructure prompted us to investigate the biogenesis of basal bodies throughout male gametogenesis using serial block face scanning electron microscopy (SBF-SEM). This method enables the reconstruction and segmentation of whole individual microgametocytes, free microgametes and female macrogametes in this mixed population of cells (see methods for details). In total 10 SBF-SEM datasets were collected consisting of at least 300 slices per dataset. Activation of gametogenesis is not synchronised, so 6 and 15 minutes post-activation samples were chosen for SBF-SEM and analysis to capture early and later events of basal body biogenesis in the microgametocyte, exflagellation events and fertilisation events.

Capturing whole microgametocytes in the early stages of basal body biogenesis via electron microscopy is challenging due the speed of early events. Male gametogenesis is a rapid process, from initial basal body formation through three rounds of mitosis to exflagellation within approximately 15 minutes. We searched for whole microgametocytes in our datasets with clustered basal bodies to identify the earliest stages when basal bodies are forming. In 300 whole microgametocytes examined, we discovered 3 microgametocytes which had 8 basal bodies clustered together. Whilst the detailed microtubule ultrastructure of basal bodies could not be resolved using SBF-SEM due to its limited resolution, each basal body was already assembling an axoneme, confirming these are basal bodies. This also confirms that there is little to no delay between basal body assembly and the initiation of axoneme assembly (Figure 2C; asterisks – basal bodies, Movie 2).

**Figure 2.**
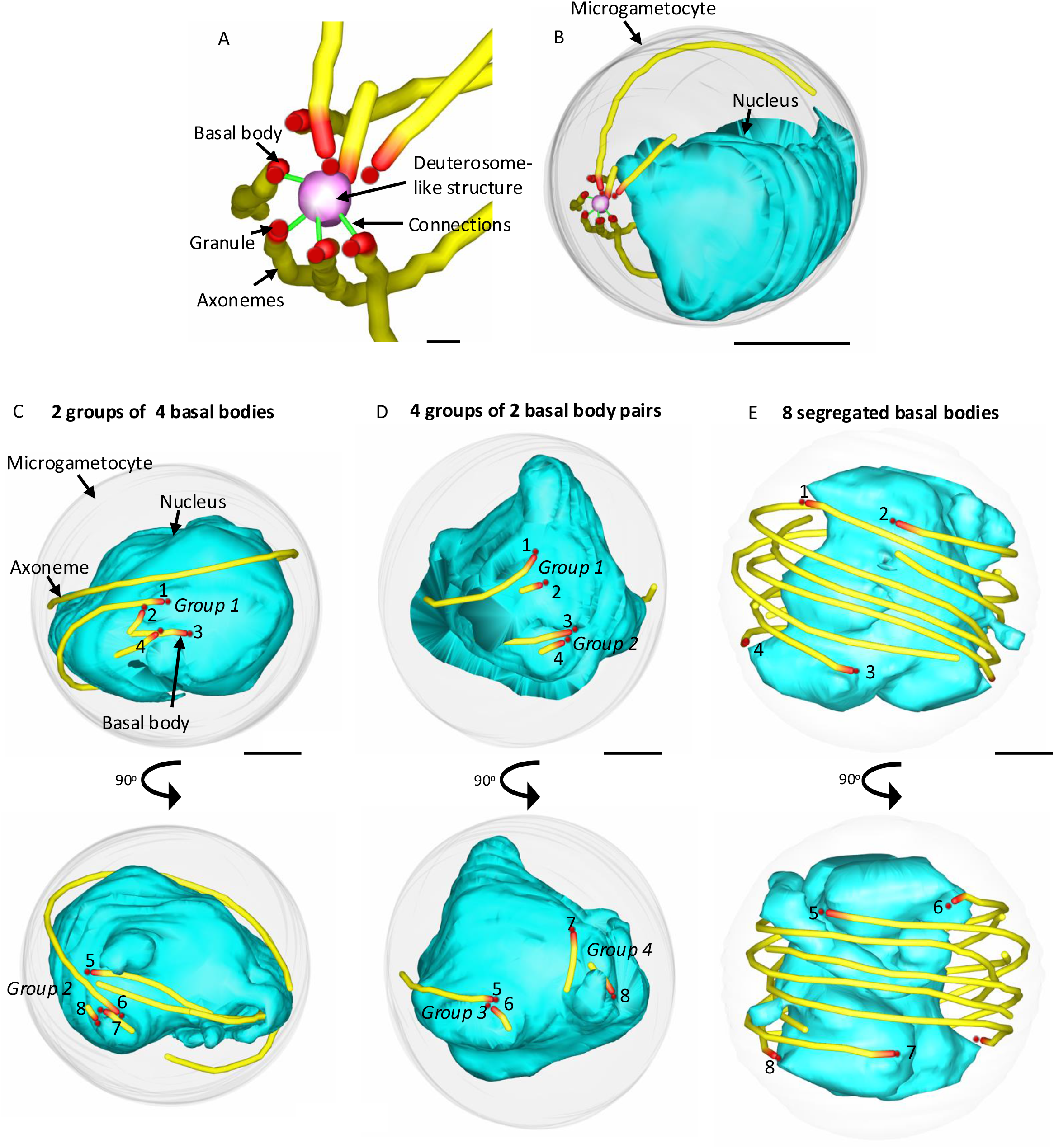
8 basal bodies surround a central electron dense structure. **A:** SBF-SEM 3D segmentation of 8 basal bodies and associated granule (red) and 8 axonemes assembling (yellow) surrounding an electron dense deuterosome-like structure (pink) with fibrous connections to some basal bodies visible (green) Scale bar = 200 nm. **B:** SBF-SEM 3D segmentation of the complete microgametocyte and nucleus from figure 2A illustrating the clustering of 8 basal bodies located close to the nucleus with the assembling axonemes. Scale bar = 1 µm. **C-E:** 3D segmentations of whole microgametocytes illustrating progression of basal body segregation through micro-gametogenesis. **C:** 2 groups of 4 basal bodies located at opposite poles following mitosis I. Group 1 with 4 basal bodies (red) in the top panel. Bottom panel is a 90° rotation of the segmentation to reveal group 2 with 4 basal bodies. Scale bar = 1 µm. **D:** 4 groups of 2 basal body pairs following mitosis II. **E:** A whole reconstructed microgametocyte with 8 individual basal bodies following mitosis III.

Intriguingly, all 8 basal bodies in the 3 microgametocytes were clustering around an electron dense structure (Figure 2A, Supplementary Figure 1, Movie 2). The electron dense structure did not contain an axoneme and was 250 nm in diameter (n=3), larger than *P. berghei* basal bodies which measured 200 nm diameter (n=24). Since all 8 basal bodies were identified in these 3 whole microgametocyte reconstructions, this electron density is very unlikely to be a basal body.

In mammalian cells there is a specialised platform for the nucleation of multiple basal bodies during multi-ciliogenesis called a deuterosome (Anderson and Brenner, 1971; Dirksen, 1971). The central spherical structure observed here with basal bodies surrounding it could be functioning as a deuterosome-like platform (Figure 2A). Fibrous connections were observed between the deuterosome-like density to some of the 8 basal bodies, suggesting a physical connection between them (Figure 2A, 2C; connections – green lines). Movie 2 shows one of the reconstructed whole microgametocytes demonstrating 8 basal bodies with assembling axonemes and the deuterosome-like density. This data confirms that all 8 basal bodies are formed at the start of gametogenesis and are organised together with their axonemes around a central deuterosome-like platform.

Bioinformatics analysis was carried out to determine if any known mammalian deuterosome proteins were conserved in *Plasmodium sp* (see methods for details). From the known human deuterosome proteins, DEUP1 (also called CCDC67), CEP63, CEP152 and CDC20, a *Plasmodium falciparum* ortholog of CDC20 could be identified by protein sequence or predicted protein structure search. BLASTp and tBLASTn searches for DEUP1, CEP63 and CEP152 identified no *Plasmodium* proteins. JACKHMMER sequence and Foldseek predicted structure searches identify some *Plasmodium* proteins, but at expectation values much higher (lower confidence) than human proteins with unrelated function like golgins, and myosins. DEUP1, CEP63 and CEP152 are predicted to have an entirely extended alpha helical structure, which contains little structural information and tends to have weak sequence conservation. *Plasmodium* likely lacks orthologs of these deuterosome proteins. There is a possibility that *Plasmodium* orthologs have diverged in sequence so far as to be undetectable, but we consider this unlikely.

Previous work in *P. berghei* demonstrated that precisely defined segregation events of basal bodies and axonemes occur during the three rounds of mitosis. During the first round of mitosis, 2 groups of 4 basal bodies are located opposite each other, this is followed by a further segregation to 4 groups of 2 pairs of basal bodies during the second mitosis and finally 8 segregated basal bodies following the third round of mitosis (Zeeshan *et al*., 2019, 2020, 2022). Whole microgametocytes were identified with 2 groups of 4 clustered basal bodies with axonemes that were located opposite to each other. The deuterosome-like platform was absent from each of these 2 clusters, suggesting that it is only required for initial basal body *de novo* assembly, but disassembles following basal body assembly (Figure 2C). Disassembly of deuterosomes also occurs in mammalian cells following assembly of basal bodies, adding further support to our analysis (Yan, Zhao and Zhu, 2016). Despite this, each group of 4 basal bodies were closely associated, suggesting there could be additional connections between basal bodies not yet identified. Further microgametocytes were identified with 4 groups of 2 closely associated basal bodies and axonemes (Figure 2D) and finally, 8 individual segregated basal bodies and axonemes (Figure 2E).

To support the basal body biogenesis stages observed by SBF-SEM, microgametocytes expressing the well-characterised basal body protein SAS6-GFP were analysed by ultrastructure expansion microscopy (U-ExM) along with centrosome marker Centrin using expansion microscopy (Figure 3 A-D) in particular to determine if all 8 basal bodies are formed at the start of gametogenesis. Timepoints post-activation of 1, 2, 4 and 8 mins were chosen to give a better chance of capturing the different stages of micro-gametogenesis. At 1 min post-activation, cells with 8 distinct SAS6–GFP foci were observed, clustered around 4 Centrin foci (Figure 3A). This confirms that all 8 basal bodies are formed at the beginning of micro-gametogenesis. Further segregation events of 2 groups of 4, 4 groups of 2 and 8 segregated basal bodies during these timepoints confirmed our recent live cell finding findings (Zeeshan *et al*., 2019, 2022; Rashpa and Brochet, 2022).

**Figure 3.**
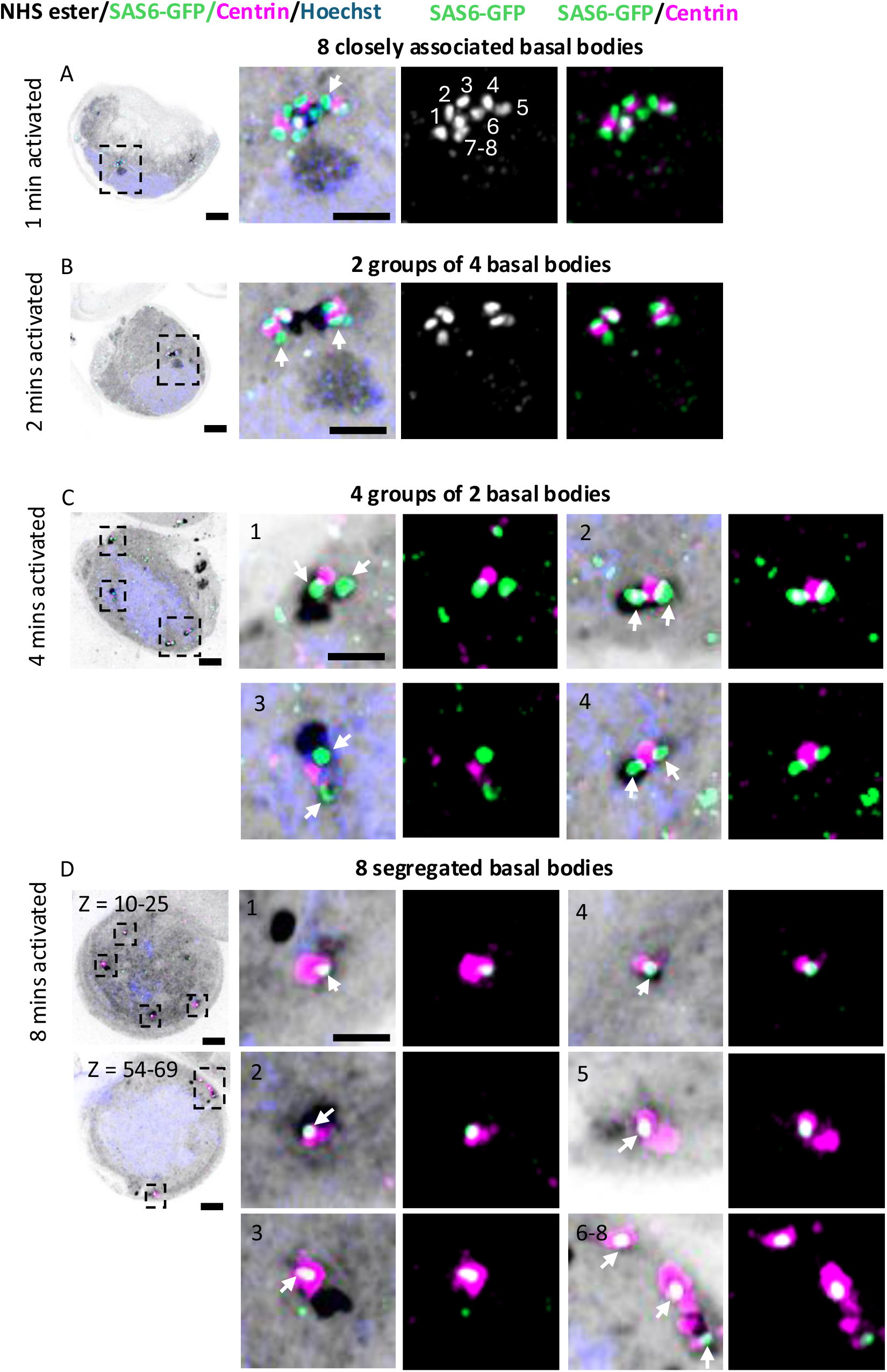
Ultrastructure expansion microscopy (U-ExM) reveals eight distinct SAS6-GFP foci following initial microgametocyte activation. **A-D:** U-ExM of whole microgametocytes showing the localisation of SAS6-GFP tagged basal bodies at **A**: 1 minute, **B:** 2 minutes, **C:** 4 minutes and **D**: 8 minutes post activation. Representative images from two independent biological experiments (n ≥ 5 cells examined per time point) are shown. **A**: 8 individual SAS6-GFP foci and 4 Centrin foci clustered together (arrows) at 1 minute post activation. **B**: 2 segregated groups of 4 SAS6-GFP (arrow) and 2 Centrin foci at 2 minutes post activation. **C:** 4 segregated groups of 2 SAS6-GFP pairs (arrows) at corners of the microgametocyte nucleus (blue). Each SAS6-GFP pair is associated with a Centrin foci. **D:** 8 individual SAS6-GFP foci (arrows), each associated with a Centrin foci at 8 minutes post activation. Images in A–C are maximum intensity projections generated from a range of z-slices encompassing all SAS6-GFP foci. For D, the annotations z = 10–25 and z = 54–69 indicate the specific z-slice ranges used to generate the maximum intensity projections. Dashed boxes indicate magnified views of the SAS6-GFP tagged basal body (arrows) and Centrin foci (pink). Scale bars = 5 µm for whole-cell images and Scale 2 µm for magnified views.

### The basal body associated granule is present during all stages of basal body biogenesis

Next, we wanted to establish if the basal body-associated granule described in Figure 1 also was assembled when basal bodies formed or if this structure was added to the proximal end of basal bodies at a later point during gametogenesis. All 300 whole microgametocytes captured by SBF-SEM were analysed for the presence of a granule associated with each basal body and an associated granule was observed in every basal body examined. The granule was present from initial *de novo* assembly of 8 basal bodies surrounding the deuterosome-like structure through to final segregation to 8 individual basal bodies at the third mitotic event (Figure 4A-C). 80 of 300 microgametocytes were undergoing exflagellation. In these cells, the granule was clearly visible as microgamete were in the process of exiting the microgametocytes (Figure 4D-E).

**Figure 4.**
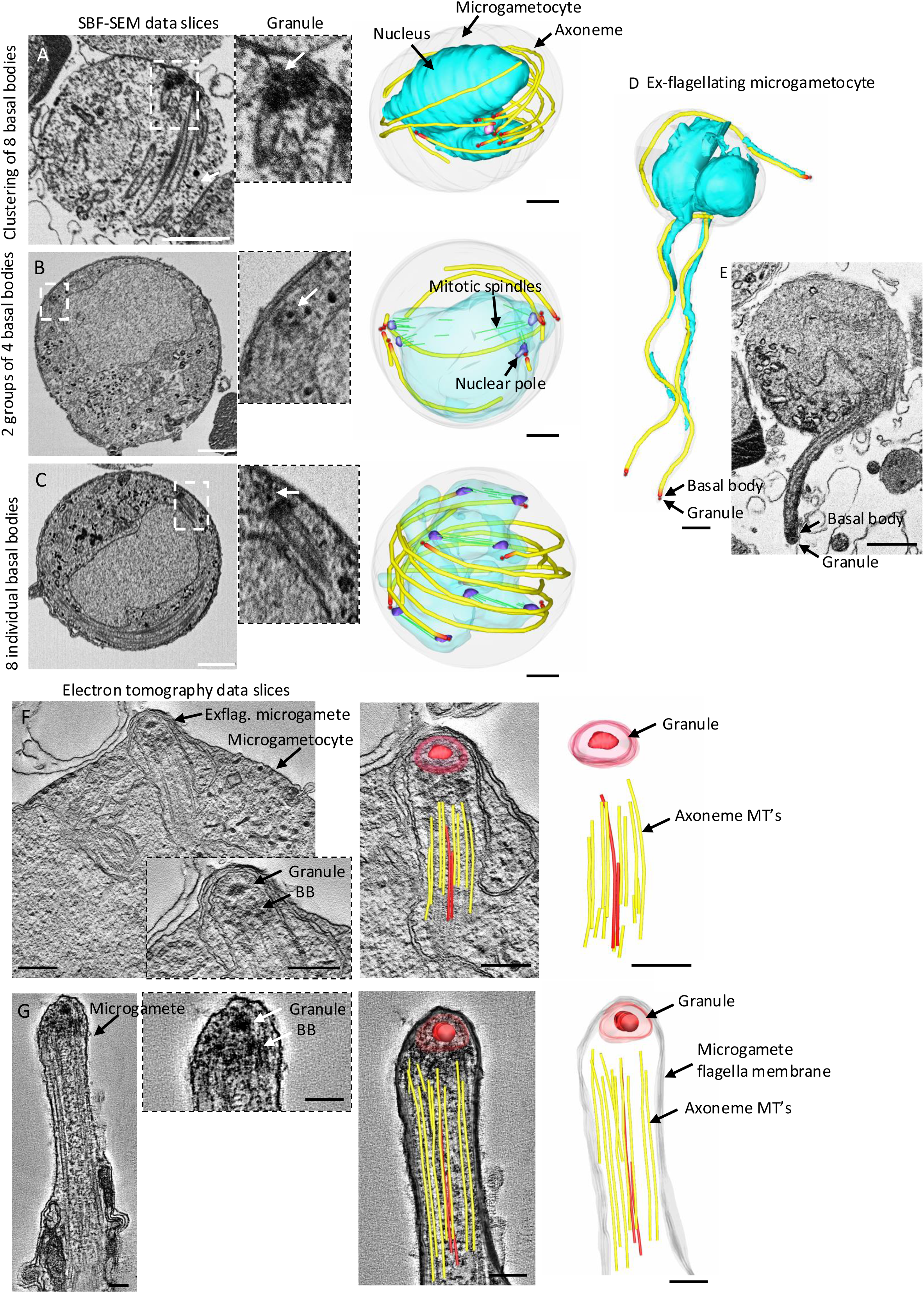
The basal body associated granule is present throughout gametogenesis and in free microgametes. **A**: SBF-SEM data slices and 3D segmentation of a microgametocyte showing a clustering of 8 basal bodies (red) around a deuterosome-like structure (pink) and the presence of the granule (SBF-SEM data slice inset – granule labelled with white arrow). Scale bar = 1 µm. Dashed box showed magnified view of the associated granule. **B:** SBF-SEM data slices and 3D segmentation of a microgametocyte showing 8 segregated basal bodies (red) into two groups of 4, mitotic spindles (green) present across the microgametocyte nucleus (cyan) and the presence of the granule (SBF-SEM data slice inset – granule labelled with white arrow). Scale bar = 1 µm. Dashed box showed magnified view of the associated granule. **C:** SBF-SEM data slices and 3D segmentation of a microgametocyte showing 8 individual basal bodies and the presence of the granule (SBF-SEM data slice inset – granule labelled with white arrow). Scale bar = 1 µm. Dashed box showed magnified view of the associated granule. **D:** SBF-SEM 3D segmentation of an exflagellating microgametocyte showing basal body and associated granule. Scale bar = 1 µm. **E:** SBF-SEM dataset of an exflagellating microgametocyte showing basal body and associated granule. Scale bar = 1 µm**. F:** electron tomography slice and 3D segmentation of an ex-flagellating microgamete emerging from a microgametocyte. Inset highlights the granule proximal to the basal body (BB), with the granule leading the microgamete out of the microgametocyte. 3D segmentation overlaid with the tomogram slice and 3D segmentation shows the budding microgamete granule, the axoneme outer doublets (yellow) microtubules (MT’s) and the central pair (red) microtubules. Scale bar = 200 nm. **G:** Tomogram slice and 3D segmentation of a free microgamete. Inset highlights the granule proximal to the basal body (BB). 3D segmentation overlaid with the tomogram slice shows the microgamete granule, the axoneme outer doublets (yellow) microtubules (MT’s) and the central pair (red) microtubules. Scale bar = 200 nm.

Serial section electron tomography was used for greater resolution of the granule close to the microgametocyte plasma membrane (Figure 4F) where the granule was clearly visible as the most proximal structure. The granule was also present in free microgametes (Figure 4G). Our findings confirm that the basal body associated granule is a permanent structure during microgamete formation.

### Microgamete-Macrogamete association is by the basal body end during fertilisation

SBF-SEM datasets contained a mixed population of whole individual microgametocytes, free microgametes and macrogametes, allowing analysis of how microgametes were interacting with female macrogametes. We identified 30 whole macrogametes with a microgamete lying across and wrapping around the surface membrane of the macrogamete (Figure 5A, Movie 3). In addition, 3 macrogametes were found to have a microgamete mid-entry into the macrogamete (Figure 5B-C, Supplementary Figure 5, Movie 4). This small number would be expected, given that entry into the macrogamete is an extremely fast process. In each case, the basal body end of the microgamete was located within the macrogamete, whilst the microgamete distal end was located outside the macrogamete (Figure 5B-C). Furthermore, the flagellar membrane was only present on the external portion of the microgamete and was not present on the internal portion (Figure 5B, C, Movies 4 and 5). This confirms that entry into the macrogamete involves fusion events of the microgamete membrane with the macrogamete membranes during entry. However, mechanistically this fusion event appears to be a directed event at a single point of entry with the basal body and granule leading the entry and fusion events occurring from the proximal end of the microgamete towards the distal end (Figure 5D). In one of the examples, the microgamete is mid-entry into a macrogamete but has not completed exflagellation from a microgametocyte (Movie 5).

**Figure 5.**
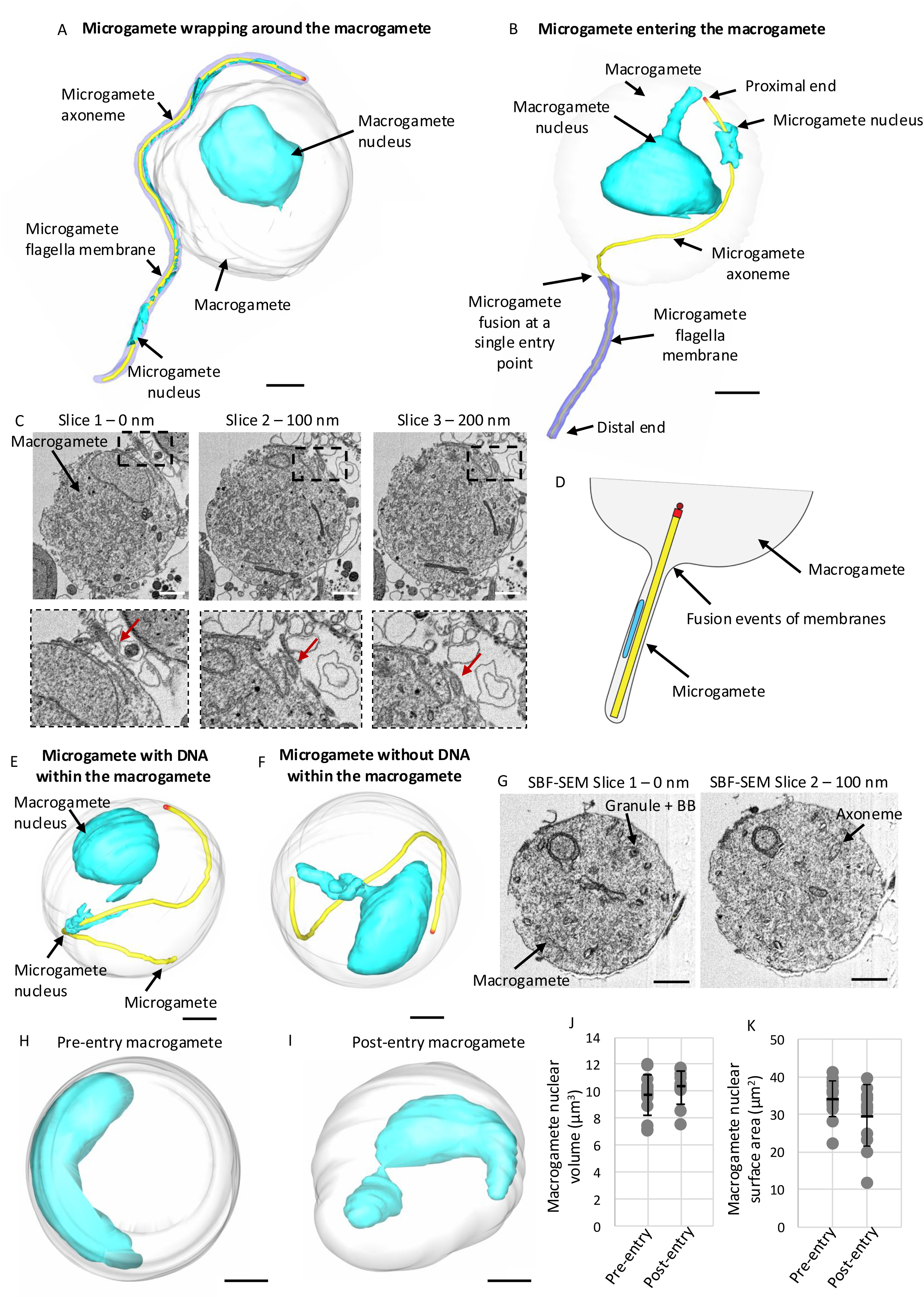
The microgamete penetrates the macrogamete basal body and granule leading entry. **A:** SBF-SEM 3D segmentation of a microgamete wrapping around a microgamete. Scale bar = 1 µm. **B:** SBF-SEM 3D segmentation of a microgamete penetrating a macrogamete. The distal end of the microgamete is outside of the microgamete and still has the microgamete flagella membrane present (purple), whist the proximal end of the microgamete is inside of the macrogamete and no longer contains the flagella membrane. Scale bar = 1 µm. **C:** SBF-SEM data slices following the microgamete (inset – red arrow) entering into the macrogamete. Scale bar = 1 µm. **D:** cartoon illustrating the single entry of a microgamete into a macrogamete. **E:** SBF-SEM 3D segmentation of a microgamete fully entered a macrogamete, with the microgamete DNA still in close association with the microgamete axoneme. Scale bar = 1 µm. **F:** SBF-SEM 3D segmentation of a microgamete fully entered a macrogamete, without a microgamete DNA present. Scale bar = 1 µm. **G:** SBF-SEM data slices of a macrogamete with the microgamete granule, basal body (BB) and axoneme present inside. Scale bar = 1 µm. **H-I**: pre-entry and post entry nucleus changes (SBF-SEM). **J:** Dot plot graph of macrogamete nuclear volume changes between unfertilised macrogametes (n=15) and macrogametes with a microgamete fully inside (n=12). **K:** Dot plot graph of macrogamete nuclear surface area changes between unfertilised macrogametes (n=15) and macrogametes with a microgamete fully inside (n=12).

Next, we searched for microgametes that had completed entry into a macrogamete (Figure 5E-G). In our datasets we identified 20 whole macrogametes with a single microgamete fully inside. These still contained an assembled 9 + 2 axoneme and a basal body with associated granule. A few microgametes still contained an associated nucleus (Figure 5E, n=3), whilst most other examples there was no associated DNA (Figure 5F, n=17). Importantly, every macrogamete contained only one microgamete inside, consistent with the presence of a mechanism that prevents the entry of additional microgametes following fertilisation, as observed in other organisms (Tsaadon *et al*., 2006; Cheeseman *et al*., 2016).

Finally, changes to the macrogamete nucleus volume and surface area were measured to investigate if there were morphological changes occurring before and after microgamete entry. Instead of the crescent-shaped nucleus of unfertilised macrogametes (Figure 5H, n=15), the nucleus of the macrogamete appeared more globular than crescent-shaped with nuclear projections following entry suggesting a conformational change in preparation for haploid genome fusion (Figure 5I) (Rashpa and Brochet, 2022; Yanase *et al*., 2026). Despite this morphological difference, no significant differences were observed in nuclear volumes (T-test, *p*=0.1595, n=27) or surface area (T-test, *p*=0.0993, n=27) between the non-penetrated macrogamete nucleus and those with a microgamete inside (Figure 5J-K).

### *HAP2* knockout microgametes can associate with the macrogamete but cannot enter

Previous work established that *HAP2* knockout microgametes were able to adhere to macrogametes but failed to fuse and fertilise, resulting no ookinete formation, thus halting progression of the life cycle (Liu *et al*., 2008; Feng *et al*., 2021; Kumar *et al*., 2022). However, a detailed high-resolution investigation of *HAP2* mutant microgametes and their association with macrogametes have not been carried out.

We used a *P. berghei HAP2* knockout mutant gametocyte activated for 30 minutes (Liu *et al*., 2008) to analyse microgamete ultrastructure and association with macrogametes using SBF-SEM. SBF-SEM datasets were searched for microgametes associated with macrogametes in the absence of *HAP2*. In total 25 whole individual macrogametes were found with one or more *HAP2ko* microgametes associating with the macrogamete plasma membrane (Figure 6A, B). There was no observable difference in the total portion of the microgamete that adhered with at least half of the microgamete length associating with the macrogamete plasma membrane. Whilst light microscopy studies have shown previously that *HAP2* microgametes are motile (Liu *et al*., 2008) and appeared normal, we carried out further ultrastructural analysis to investigate any observable changes. Microgametes still contained a basal body and basal body associated granule when located within a microgametocyte during gametogenesis (Figure 6C) and in free microgametes (Figure 6D). Nine-fold rotational averaging analysis was used to investigate the ultrastructural organisation of the 9+2 axoneme. *HAP2* and wildtype 9+2 axonemes demonstrated no visible defects in axoneme organisation with the ninefold outer doublet symmetry and central pair unchanged (Figure 6E, n=20 per group). These results show that there are no observable defects in the *HAP2* microgamete ultrastructure and confirms that *HAP2* microgametes adhere to macrogametes but fail to initiate basal body directed entry.

**Figure 6.**
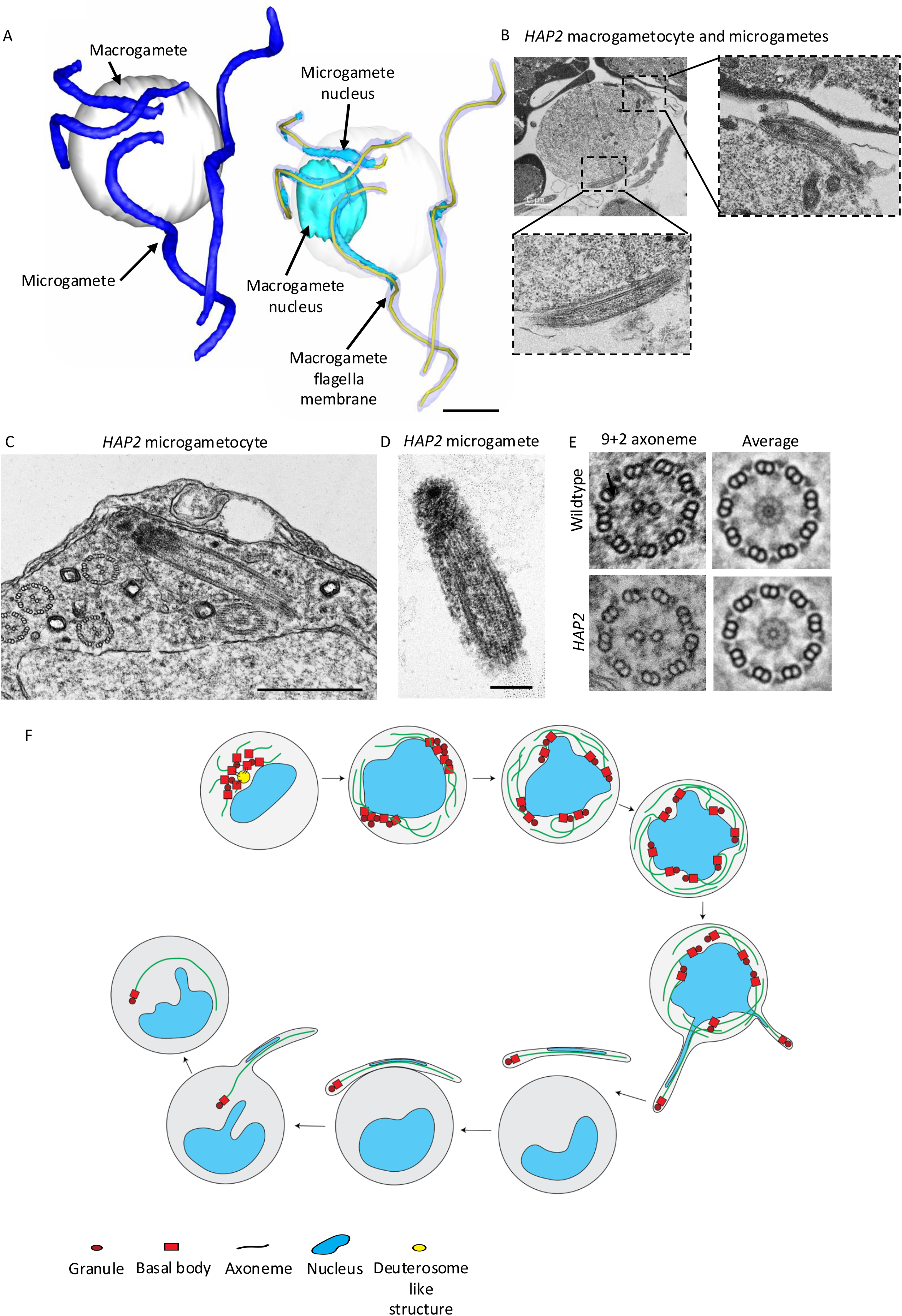
*HAP2* microgametes still associate with the macrogamete despite entry blocked. **A:** SBF-SEM 3D segmentation of four *HAP2* microgametes making close contract with a macrogamete but no penetration was observed. Scale bar = 1 µm. **B:** TEM micrographs of two *HAP2* microgametes making close contact with a macrogamete. **C:** TEM micrograph of a *HAP2* microgametocyte showing the presence of the basal body, granule and axoneme. Scale bar = 500 nm. **D:** TEM micrograph of a *HAP2* microgamete showing the presence of the basal body, granule and axoneme. Scale bar = 200 nm. **E:** Ninefold rotational averaging of wildtype and *HAP2* microgametocyte axonemes (n=20 per group). **F**: Schematic representation of stages of basal body biogenesis and fertilisation; 1. Activated microgametocytes form 8 basal bodies and associated granules *de novo* clustering around a deuterosome-like structure and axonemes assembly begins; 2. Basal bodies and associated granules segregate into 2 groups of 4, axoneme elongation continues and DNA replication begins (2N); 3. Basal bodies segregate into 4 groups of 2, axoneme elongate and DNA replication continues (4N); 4. Basal bodies segregate and DNA replication continues (8N); 5. Exflagellation occurs with the basal body associated granule leading to form a microgamete; 6. Free motile microgamete and unfertilised macrogamete; 7. Microgamete contacts macrogamete plasma membrane; 8. Microgamete penetrates macrogamete and fusion of microgamete and macrogamete occurs at a single-entry point; 9. Microgamete fully penetrated macrogamete for fertilisation.

## Discussion

Using volume electron microscopy (SBF-SEM) and serial section electron tomography (ssET), we have enhanced our understanding and knowledge of the *P. berghei* de novo basal body ultrastructure from initial assembly to fertilisation (Figure 6F). Our data revealed that the eight basal bodies formed *de novo* following gametocyte activation were clustered around and closely associated with a previously uncharacterised structure we term deuterosome-like. Formation of multi-ciliated cells in mammals often requires a deuterosome, which functions as a platform for *de novo* basal body nucleation, with multiple deuterosomes producing hundreds of basal bodies in a single differentiated cell. The basal bodies then migrate and dock to the apical surface and produce cilia (Kim and Dynlacht, 2013; Mercey *et al*., 2019). In multi-ciliated mammalian cells, deuterosomes are present during assembly of basal bodies before disappearing once amplification is complete. This is the same pattern of presence and absence that we observed for the *P. berghei* deuterosome-like structure where it was only present during early stages of gametogenesis. The deuterosome is not well characterised at a molecular level and only three proteins DEUP1 (also called CCDC67), CEP63, CEP152 and CDC20 have been identified in mammalian multi-ciliated cell deuterosomes (Zhao *et al*., 2013; Boutin *et al*., 2025; Lee *et al*., 2026). Our bioinformatics analysis identified only CDC20 - an anaphase-promoting complex protein which has been found to be essential for male gametogenesis but is not essential for mitosis in the asexual blood stage. in *P. berghei.* Using serial section electron tomography, we have resolved the ultrastructure of the *P. berghei* basal body and demonstrated that it is an unusually short basal body at only ∼50nm compared to many other organisms, which are ∼200nm or more and only composed of 9 single microtubules rather than the usual triplet microtubule configuration (Allen, 1969; Dutcher and O’Toole, 2016; Tassin, Lemullois and Aubusson Fleury, 2016; Vaughan and Gull, 2016). We also characterised a basal body associated granule that was present at the proximal end of the basal body. It was observed during the initial formation of 8 basal bodies when they are associated with the deuterosome-like structure and was still present following entry into the macrogamete during fertilisation. Whilst its precise function and molecular content is currently unknown, its location at the proximal end of the basal body could point to functions in docking of the axoneme of the immature microgamete to the microgametocyte plasma membrane when exflagellation occurs and/or it could be important for initial entry into the macrogamete like the acrosome of the mammalian sperm during fertilisation (Tsaadon *et al*., 2006; Berruti and Paiardi, 2011). This granule was first discovered in *Plasmodium yoelii* microgametocytes suggesting it is a conserved structure among *Plasmodium species* (Sinden, Canning and Spain, 1976), but further work will be required to identify molecules to study the function of this intriguing structure.

We reveal that, although the microgametes associated with the macrogamete membrane, fusion and entry occurs via a single point on the macrogamete with the basal body end leading the entry. Following *HAP2* knockdown, we found that the basal body associated granule remained and were no observable ultrastructural defects. The inability of the microgametes to penetrate the macrogametes is consistent with a defect in membrane fusion rather than in gamete recognition or structure. Indeed, the basal body granule could define a specific membrane domain that is important for initial events in fusion and/or entry. Previous work has shown that there is a two-step process in microgamete fertilisation. The first is microgamete adhesion with adhesion molecules such as p48/45 required for this (Alkema *et al*., 2024). The second is microgamete fusion with the macrogamete membrane which is known to be *HAP2-*mediated. Future studies to identify a basal body proteome and its essential interacting partners may help to decipher the molecules associated with granule.

## Methods

### Ethics statement

All animal experiments were conducted in the United Kingdom in accordance with the Animals (Scientific Procedures) Act 1986 and approved by the Home Office under Project Licence numbers PDD2D5182 and PP3589958. Protocols were reviewed and approved by the institutional Animal Welfare and Ethical Review Body (AWERB) prior to implementation. Procedures were designed to minimise animal suffering and the number of animals used, in line with the 3Rs (Replacement, Reduction, Refinement) principles. Experiments were performed on female CD1 outbred mice aged 6–8 weeks, housed under standard conditions with environmental enrichment and monitored daily for health and welfare. The conditions of mice kept are a 12 h light and 12 h dark (7 till 7) light cycle, the room temperature is kept between 20 and 24 °C and the humidity is kept between 40 and 60%.

### Parasite culture and gametocyte purification

*Plasmodium berghei* transgenic lines were injected into phenylhydrazine-treated mice (Beetsma *et al*., 1998). Gametocyte enrichment was achieved by sulfadiazine treatment after 2 days post-infection. The blood was collected on day 4 post-infection, and gametocyte-infected cells were purified on a 48% v/v NycoDenz gradient. The 48% NycoDenz solution was prepared by diluting a NycoDenz stock solution (27.6% w/v NycoDenz in 5 mM Tris-HCl, pH 7.20, 3 mM KCl, 0.3 mM EDTA) with coelenterazine loading buffer (CLB; PBS containing 20 mM HEPES, 20 mM glucose, 4 mM sodium bicarbonate, 1 mM EGTA, and 0.1% w/v bovine serum albumin, pH 7.25). The gametocytes were harvested from the interface and activated in ookinete culture medium (RPMI 1640 medium containing 25 mM HEPES, pH 7.5, 10% FCS, and 100 μM xanthurenic acid).

### Transmission electron microscopy (TEM)

For ultrastructural analysis of gametocytes and gametes, cells were fixed in 4% glutaraldehyde in 0.1 M phosphate buffer at 6, 15 and 30 minutes post activation. Samples were processed for transmission electron microscopy. Briefly, fixed cells were post-fixed in osmium tetroxide, stained *en bloc* with uranyl acetate, dehydrated through a graded ethanol series, and embedded in Spurr’s epoxy resin. Ultrathin sections (70 nm) were cut using a diamond knife, mounted on copper mesh grids, and post stained with uranyl acetate and lead citrate. Sections were examined using a Joel JEM 1400 Flash TEM at 120 kV.

### Serial block face scanning electron microscopy (SBF-SEM) of culture derived gametocytes

Fractions enriched in *Plasmodium berghei* gametocytes (produced as described above) were fixed at room temperature in 2.5% glutaraldehyde in 0.1 M phosphate buffer, 6, 15 and 30 minutes after activation, to allow for an asynchronous population containing the different stages of gametogenesis and gametes to be analysed. Samples were then spun and washed three times in 0.1 M phosphate buffer, and post-fixed in 1% osmium tetroxide in 1.5% potassium ferrocyanide in 0.1 M phosphate buffer (for 45 min at room temperature, and in the dark). After the first osmium step, samples were washed three times in 0.1 M phosphate buffer, incubated in 1% tannic acid in 0.1 M phosphate buffer for 30 min at room temperature, and then subjected to a second osmium step (2% OsO_4_ in ddH_2_O, for 30 min, at room temperature, and in the dark). Samples were then incubated in 2% uranyl acetate in ddH_2_O for 2 h, dehydrated in acetone (a progressive series of acetone concentrations from 20%, 40%, 90% to 100% with three changes in molecular-sieved ultradry acetone over 4 h) and embedded in TAAB 812 Hard resin (TAAB, catalogue number T030). The tips of resin blocks containing samples were trimmed and mounted onto aluminium pins using conductive epoxy glue and silver dag and then sputter coated with a layer (10–13 nm) of gold, in an Agar Auto Sputter Coater (Agar Scientific). Before SBF-SEM imaging, ultrathin sections (70 nm) of the block face were examined in a Jeol JEM 1400 Flash transmission electron microscope (JEOL), to verify sample quality. Samples were then imaged in a Merlin VP compact high resolution scanning electron microscope (Zeiss) equipped with a 3View stage (Gatan-Ametek), and an OnPoint back-scattered electron detector (Gatan-Ametek), in variable pressure. The following imaging conditions were used: 2 kV, 30 µm aperture, 100% focal-charge compensation, 2 nm pixel size, 5 μs pixel time, 100 nm section thickness.

### Dual axis cellular electron tomography

Fractions enriched in *Plasmodium berghei* gametocytes (produced as described above) were fixed in 4% glutaraldehyde in 0.1 M phosphate buffer and processed for electron microscopy. Briefly, samples were post-fixed in osmium tetroxide, treated en bloc with uranyl acetate, dehydrated in acetone and embedded in Spurr’s epoxy resin. Thin sections were stained with uranyl acetate and Reynolds’ lead citrate prior to examination in a JEOL JEM-1400 Flash transmission electron microscope (JOEL, UK). Serial-section cellular electron tomography (ssET) was performed on 150 nm sections collected onto formvar-coated slot grids. Grids were mounted in a Fishione dual-axis tomography holder (Fischione instruments) and dual-axis tilt-series (55° to −55°, with 1° tilt between images) of the same area in consecutive sections were acquired using SerialEM (Mastronarde, 2005). Tomogram generation and serial tomogram joining were performed in ETomo (Kremer, Mastronarde and McIntosh, 1996).

### SBF-SEM and tomography data segmentation and analysis

Data were processed using the IMOD software package (Kremer, Mastronarde and McIntosh, 1996). Image stacks were assembled, corrected (for z scaling and orientation) and aligned using Etomo, and 3D segmentation was produced using 3dmod (Kremer, Mastronarde and McIntosh, 1996). Whole individual cells were identified and manually segmented from trimmed regions of original datasets. The microgametocytes, macrogametocytes and microgametes analysed represented all whole individual cells of each type found in the SBF-SEM datasets. In each cell, the cell membrane, the nucleus, the axonemes with subtending basal bodies and the nuclear poles (microgametocyte only) were identified based on distinctive ultrastructural features and then segmented manually as individual objects (Hair *et al*., 2023). Axonemes were identified by the outer microtubule doublets as an electron dense circle with the central pair appearing as a central electron density. Basal bodies were segmented as an electron dense structure at the proximal end of the axoneme. Nuclear poles were segmented as an electron dense structure positioned within the nucleus close to the nuclear membrane. Volume and surface area for the nucleus were obtained in 3dmod, based on objects surface rendering. Statistical analysis of macrogametocyte nucleus volume and surface area (T-test) were calculated in Microsoft Excel.

### Ninefold rotational averaging of *P. berghei* axonemes

For generation of averaged axoneme views, doublet A tubule centres in axoneme images were identified, fitted to an ellipse then perspective corrected to ensure circularity, followed by ninefold rotational averaging (Gadelha, Cunha-e-Silva and de Souza, 2013). Fifteen rotationally averaged axonemes were then aligned and averaged. Difference maps were generated by comparison with the average 15 rotationally averaged parental cell line, and per-pixel statistical significance of electron density changes calculated by Mann Whitney U test (with multiple-comparison correction for the number of pixels within the axoneme cross-section) (Gray, Fort and Wheeler, 2024).

### Generation of transgenic parasites

For C-terminal GFP-tagging of SAS6 (PBANKA_0106200) by single crossover homologous recombination, the 3’ region immediately upstream of the stop codon (omitting the stop codon) was amplified using primers (T1891: 5’-CCCCGGTACCGCATAGAATGTGAAAAATGTAAACTGG-3’ and T1892: 5’-CCCCGGGCCCAATTCCGGGAGGAATAAATTTCACG-3’). The DNA fragment was inserted using KpnI and ApaI restriction sites upstream of the *gfp* sequence in the p277 plasmid containing a human *dhfr* cassette to confer resistance to pyrimethamine. *P. berghei* ANKA line 2.34 parasites were then transfected by electroporation as described previously (Janse *et al*., 2006; Guttery *et al*., 2012). A schematic representation of the endogenous gene locus, the construct, and the recombined gene locus can be found in Supplementary Figure 3. Genotype analysis was performed using integration PCR as illustrated in Supplementary Figure 3. Integration was confirmed using a forward primer upstream of the homology region (INT T189: 5’-GTAATGAATATAGTGAGGTAAATAAAAG-3’) and a *gfp*-specific reverse primer (ol 492: 5’-ACGCTGAACTTGTGGCCG-3’).

### Ultrastructure expansion microscopy (U-ExM)

U-ExM was performed on PbSAS6-GFP activated microgametocytes. Parasites were fixed in 4% formaldehyde in MTSB buffer (10 mM MES, 150 mM NaCl, 5 mM EGTA, 5 mM MgCl_2_, 5 mM glucose, pH7.0) and adhered to 10 mm poly-D-lysine–coated round coverslips for 15 min.

Coverslips were incubated overnight at 4°C in 1.4% formaldehyde (FA)/2% acrylamide (AA). Gelation was performed in ammonium persulphate/TEMED (10% each)/monomer solution (23% sodium acrylate; 10% AA; 0.1% BIS-AA in PBS) on ice for 5 min and at 37°C for 30 min. Gels were denatured for 15 min at 37°C and for 90 min at 95°C in denaturation buffer (200 mM SDS, 200 mM NaCl, 50 mM Tris, pH 9.0, in water). After denaturation, gels were incubated in distilled water overnight for complete expansion. The next day, circular gel pieces with a diameter of 13 mm were excised, and the gels were washed in PBS three times for 15 min to remove excess water. The gels were then incubated in blocking buffer (3% BSA in PBS) at room temperature for 30 min, incubated with rabbit polyclonal anti-GFP antibody (1:250 dilution; A11122; Invitrogen) and mouse monoclonal anti-centrin antibody, clone 20H5 (1:250 dilution: 04-1624; Sigma-Aldrich) in blocking buffer at 4°C overnight. The gels were washed three times for 15 min in wash buffer (0.5% v/v Tween-20 in PBS) and then incubated with 8 μg/ml ATTO 665 NHS-ester (76245, Sigma-Aldrich), 10 μg/ml Hoechst 33342 (Molecular Probes), and Alexa Fluor 488 goat anti-rabbit IgG (A11008; Invitrogen) and Alexa Fluor 568 goat anti-mouse IgG (A11004; Invitrogen) in PBS (1:500 dilution) at 37°C for 2.5 h. Blocking and all antibody incubation steps were performed with gentle shaking. The gels were then washed three times for 15 min with wash buffer and expanded overnight in ultrapure water. The expanded gel was placed in a 35-mm glass-bottom dish (MatTek) with the 14-mm glass coated with poly-D-lysine. High-resolution images were acquired on a Zeiss CellDiscoverer 7 with Airyscan using a 50×/1.2 water objective. Confocal z-stacks were acquired using line scanning and the following settings: 55 × 55 nm pixel size, 170-nm z-step, 1.07 μs/pixel dwell time, gain settings of 650 (for 405, 561 and 640 nm) and 850 (for 488 nm), and laser powers of 3.5% (405 nm), 4.5% (488 nm), 5.0% (561 nm), and 5.0% (640 nm). The z-stack images were processed and analysed using Fiji (version 1.54f). Each experiment was repeated two times, and protein localisation was assessed in 5–6 cells per condition.

### Generation of HAP2 knockout cell line

To replace all protein-coding sequence of the *HAP2* gene (GenBank accession no. XM 671808 with a *T. gondii dhfr/ts* expression cassette conveying resistance to pyrimethamine, a targeting vector was constructed in plasmid pBS-DHFR. A 736-bp fragment comprising 5′-flanking sequence immediately upstream of the start codon was amplified from *P. berghei* genomic DNA using primers GF-1 (5′-CCCC<u>GGGCCC</u>GCGCGTTATTATTATTCGGGC-3′, restriction site underlined) and GF-2 (5′-GGGG<u>AAGCTT</u>TTTTTC TAAATGAAATATTAAAGAATGGC-3′) and inserted into ApaI and HindIII restriction sites upstream of the *dhfr/ts* cassette of pBS-DHFR. A 967-bp fragment of 3′-flanking sequence was then generated using primers GF-3 (5′-CCCCG<u>AATTCA</u>T TACATGGAATAGTATTTGCAAATTTG-3′) and GF-4 (5′-GGGG<u>TCTAGA</u>CAATATACATGCTGATAACCTCC-3′) and inserted downstream from the *dhfr/ts* cassette using EcoRI and XbaI restriction sites. The replacement construct was excised as an ApaI/XbaI fragment and used for the electroporation of cultured *P. berghei* schizonts as previously described (Janse, Ramesar and Waters, 2006). Following dilution cloning of drug-resistant parasites, genotyping of two *hap2* clones was done by Southern blot hybridization on EcoRI-digested genomic DNA using the ApaI/HindIII fragment of 5′ targeting sequence as a probe. Diagnostic PCR analysis used primers GFko1 (5′-CTCGAATATGTAGATATATCCA GATG-3′) and GFko2 (5′-CAGAGATGTTATAGCTAGT GATATAAC-3′) specific for *HAP2*, and primers GFint (5′-CTAAGTAGCAACTATTTTGTAAAATTATATC-3′) and 70 (*1*) to span the predicted 5′ integration site.

### Bioinformatics analysis

Reference sequences and basic properties for human deuterosome proteins were taken from Uniprot (The UniProt Consortium, 2025): Q05D60 (DEUP1/CCDC67), Q96MT8 (CEP63), O94986 (CEP152/KIAA0912) and Q12834 (CDC20). BLASTp and tBLASTn searches were performed using PlasmoDB, part of VeuPathDB (Alvarez-Jarreta *et al*., 2024), against the predicted proteome and genome of *Plasmodium falciparum* 3D7, *Plasmodium yoelii* 17X and *Plasmodium berghei* ANKA. JACKHMMER searches used the EBI web interface (Madeira *et al*., 2022), searching with 3 iterations and otherwise default settings against Uniprot predicted proteomes. Alphafold2 (Jumper *et al*., 2021) predicted protein structures of the human proteins were taken from the EBI Alphafold database (Varadi *et al*., 2022, 2024), and the Foldseek server used for protein structure similarity searches against EBI Alphafold database predicted structures (van Kempen *et al*., 2024).

---

**Supplementary Figure 1: The presence of a deuterosome-like structure.** Sequential SBF-SEM data slices showing a microgametocyte with a clustering of 8 electron dense basal bodies (asterisks) from which axonemes elongate from, surrounding a central electron density termed deuterosome-like structure (white arrow). Filamentous connections can be seen between some of the basal bodies (green arrow). Scale bar = 1 µm.

**Supplementary Figure 2:** Two further examples of a microgamete mid-entry into a macrogamete using SBF-SEM.

**Supplementary Figure 3: Generation of PbSAS6-GFP parasites. A:** Schematic representation of the endogenous *sas6* locus, the GFP-tagging construct, and the recombined *sas6* locus following single homologous recombination. Arrows indicate the position of PCR primers used to confirm successful integration of the construct. **B:** Diagnostic PCR of *sas6* and WT-GFP parasites using the diagnostic primers to show the correct integration. Integration of the *sas6* tagging construct gives a band of 1551 bp.

**Movie 1:** Serial cellular electron tomogram of basal body and associate granule. basal body is formed of 9 single microtubules (yellow) which are each associated with an electron density (purple), a second outer doublet microtubule B-tubule (green) forms on the side of the single A-tubule distal to the basal body to form the axoneme. Central pair microtubules (green). Basal body associated granule. Central electron density – pink, filamentous material – dark pink, and an electron dense ring – red.

**Movie 2:** SBF-SEM dataset and 3D reconstruction of a whole microgametocyte with 8 basal bodies (red), with growing axonemes (yellow). The basal bodies are clustered around a deuterosome-like structure (purple), nucleus (blue).

**Movie 3:** SBF-SEM dataset and 3D reconstruction of a whole macrogamete with a single microgamete adhering. Basal body (red), axoneme (yellow), nucleus (blue).

**Movie 4 - 5:** SBF-SEM dataset and 3D reconstruction of 2 whole macrogametes with a single microgamete mid-entry into the macrogamete.

## Acknowledgements

We thank Dr Mohammed Zeeshan and Dr David G Guttery for their support and discussion. The funding for this work is supported by an ERC advance grant funded by UKRI Frontier Science (EP/X024776/1), MRC UK (MR/K011782/1), and BBSRC (BB/L013827/1, BB/X014681/1) to RT. MH and DB were supported as research fellows and senior technician on UKRI Frontier Science (EP/X024776/1). RY supported by the BBSRC (BB/X014681/1).

## Author Contributions

MH, SV and RT conceptualised the project.

MH and SV performed all the ultrastructure serial section tomography and SBF-SEM work on the *P. berghei* parasite samples provided by RT and RY.

DJPF provided the support and analysis for the TEM and SBF-SEM work.

RY and RT performed all *P. berghei* expansion microscopy, cell and molecular biology work.

DB designed and transfected the SAS6tg and provided the support with all animal work.

RW provided the bioinformatic support the deuterosome analysis.

MH, SV and RT wrote the first draft and all other authors reviewed and edited the manuscript.

