## Supplementary Figures for "Ultrastructural dynamics of basal bodies during microgamete formation and fertilisation in *Plasmodium*"

Supplementary Figure 1

Sequential slices through the basal body area from a whole microgametocyte shown in Figure 2A and Movie 2

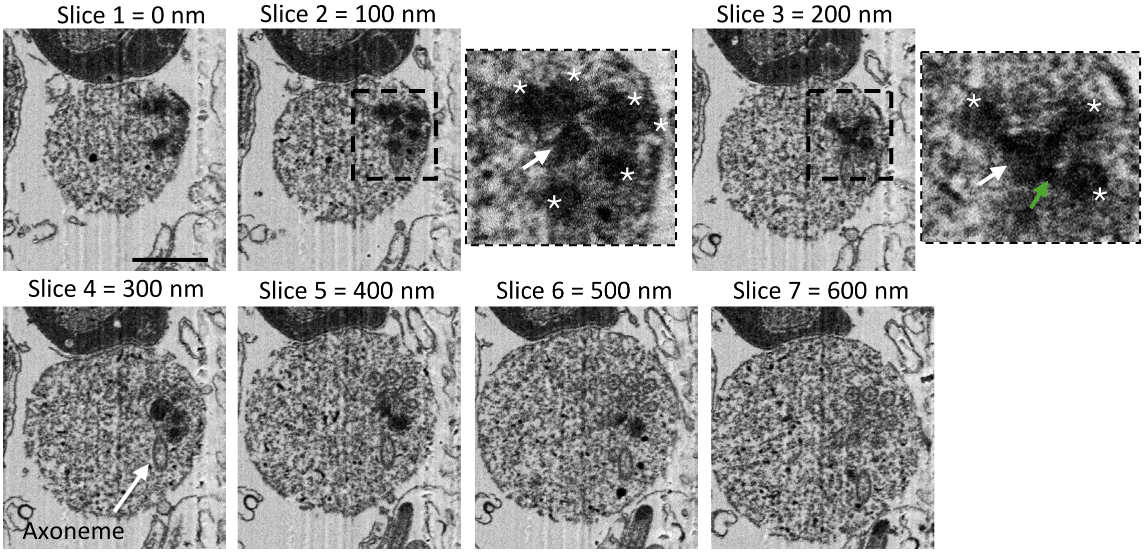

Supplementary Figure 2

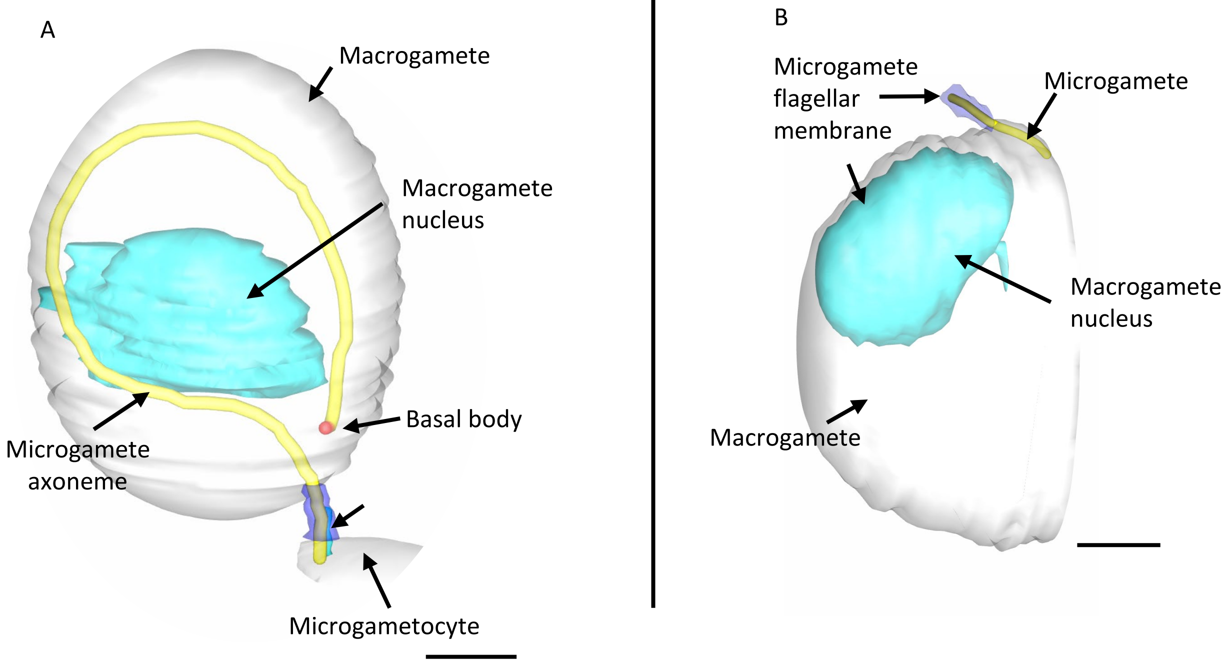

Supplementary Figure 3

A

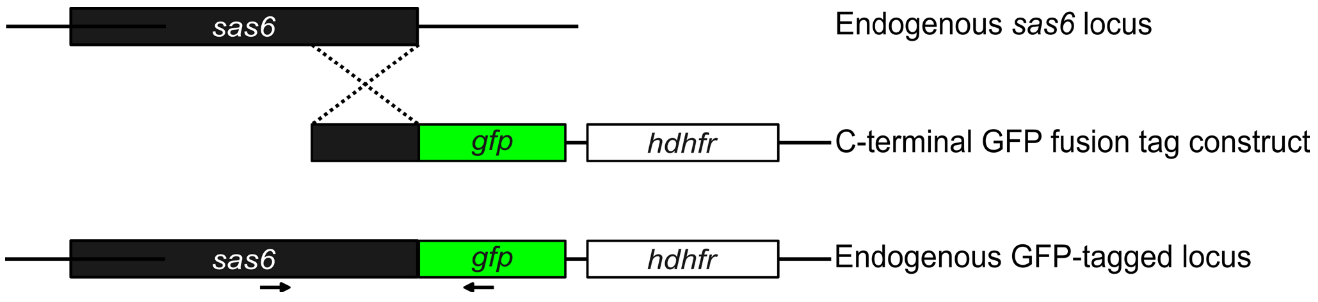


B

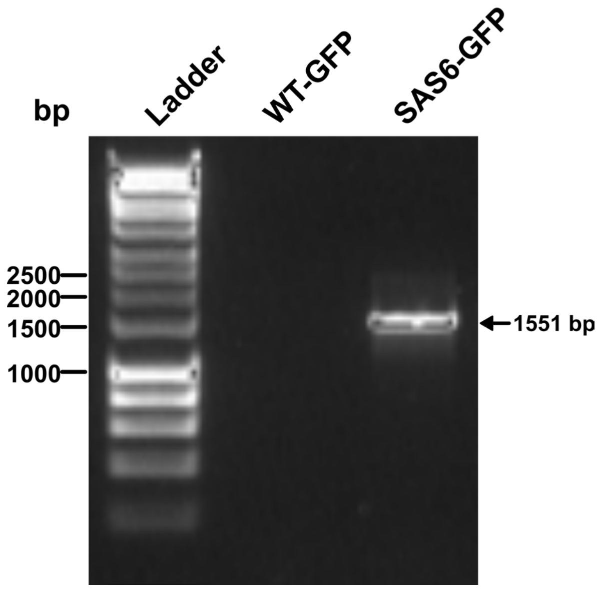
